# Does extrinsic mortality accelerate the pace of life? A bare-bones approach

**DOI:** 10.1101/777698

**Authors:** Jean-Baptiste André, François Rousset

## Abstract

It is commonly asserted that when extrinsic mortality is high, individuals should invest early in reproduction. This intuition thrives in the literature on life-history theory and human behavior, yet it has been criticized repeatedly on the basis of mathematical models. The intuition is indeed wrong; but a recent theoretical criticism has confused the reason why it is wrong, thereby obscuring earlier and sounder criticisms. In the present article, based on the simplest possible model, we sought to clarify these issues. We confirm earlier findings that extrinsic mortality can affect the evolution of pace of life, not because it leaves little time to reproduce, but through its effects on density-dependent competition. This result highlights the importance of accounting for density-dependence in theoretical models and data analyses. Further, we find little support for the recent claim that the direction of selection on a reaction norm in a variable environment cannot be easily inferred from models made in homogeneous environments. In conclusion, although life-history theory is still imperfect, it has provided simple results that deserve to be understood.

## 2 Introduction

Life history theory is widely used to interpret in an adaptive way a number of inter-individual differences observed within the human species, that can be linked to variations in individuals’ environments. In particular, consistent data show that people living in harsh environments (poor, dangerous, and/or uncertain environments; the notion of harshness being partly ambiguous) invest more, and earlier, in reproduction, and less in the growth and maintenance of their biological capital (see Pepper and Nettle 2017 for a review). In the context of life history theory, this observation is interpreted as the plastic expression of a “fast” strategy supposed to be adaptive in harsh environments.

Not all the dimensions of the physiology and behavior of people living in harsh conditions are well characterized empirically and, above all, not all have a clear adaptive explanation. Life history approaches still require many developments to account for the whole logic of human intra-specific variability (see Mathot and Frankenhuis 2018 for a review). But there is at least one intuition in this literature, that has –bibliometrically if not scientifically– survived previous discussions: natural selection should have led to the evolution of a reaction norm whereby individuals adapt their life history to the level of extrinsic mortality they perceive in their environment, investing more in reproduction and less in survival when mortality is high, and vice versa when it is low. When extrinsic mortality is high (i.e., in a harsh environment), individuals have a greater chance of dying before they have had time to reproduce and therefore, so the intuition goes, they must invest early and intensively in reproduction, to increase their chances of reproducing before they die. That is, they must choose a fast pace of life. Conversely, when extrinsic mortality is low, individuals are unlikely to die young, hence they can afford to invest in growth and survival and to delay their reproduction. That is, they can choose a slow pace of life. This intuitive understanding of the effect of extrinsic mortality was initially spelled out by Williams (1957), and is commonly recalled in the evolutionary biology literature, as well as in the literature on evolution and human behavior (see e.g., Ellis et al., 2009; Nettle, 2010; Belsky et al., 2010; Griskevicius et al., 2011; Frankenhuis et al., 2013; Mell et al., 2018).

Yet this intuitive understanding is not actually supported by evolutionary theory. Careful models show that the effect of extrinsic mortality does not always go in this direction, and more importantly that it does not occur for the intuitive reason posited in Williams’ hypothesis. Several articles, including some classics, have well demonstrated this (e.g. Abrams 1993; Williams et al. 2006; Caswell 2007; Dańko et al. 2017, 2018; Moorad et al. 2019). Unfortunately, they are little known in the literature on human behavior. To make matters worse, the only paper that has sought to communicate this notion to the community of evolutionary psychologists and evolutionary anthropologists is a preprint published on bioR*χ*iv (Baldini, 2015) that brings up the problem with Williams’ hypothesis, but also presents some claims that are not correct. Notably, Baldini (2015) is based on a definition of extrinsic mortality that differs from the definition usually used in the evolutionary literature (see also Del Giudice, 2019 in this special issue). Beyond trying to debunk Williams’ hypothesis, Baldini (2015) also questions the possibility of understanding plastic strategies in variable environnments from models of evolution in constant environments. As we shall see, his conclusions on plasticity are misleading.

Here our objective is to expose in the simplest possible way the main points that need to be understood, by scholars in evolutionary psychology and evolutionary anthropology, regarding the effect of extrinsic mortality on the evolution of pace of life, including in a plastic species living in a variable environment. Our analysis is simpler than previous works, which rested on the mathematics of age-structured populations which may obscure some simple messages. However, we concur with the conclusions of these previous works. In particular, in line with Dańko et al. (2018), we stress that the importance of taking density dependence into account is still under-appreciated in broad discussions of life-history evolution –although accounted in a number of specific models– and that too little effort is put to document pathways of density-dependence in natural populations. The same problems have been met in the study of spatially structured populations (Leturque and Rousset, 2004).

Our aim is thus to build a minimal model that captures the key findings of this literature in the simplest possible way (in particular the results of Dańko et al. 2017). In a nutshell these findings are the following. (1) Extrinsic mortality does *not* directly affect the evolution of life history traits. (2) Extrinsic mortality only affects the evolution of life history *indirectly* through its effect on the intensity of competition. A higher extrinsic mortality reduces the intensity of competition whereas a lower mortality leads to more competition. (3) This modification of the intensity of competition can affect in turn the evolution of life history and this effect can be understood in a principled manner (4) *Quantitatively* speaking, the effect of extrinsic mortality on life history in a plastic species living in a variable environment may be different from its effect in a hard-wired species living in a constant environment. (4) But, *qualitatively*, the directions of the effects are the same in both cases.

## 3 Modelling approach

### 3.1 Measuring the effect of selection in a homogeneous environment

The objective of the following analysis is to measure the effect of natural selection on a quantitative trait that controls the pace of life of individuals, or more generally a quantitative trait that affects both their fecundity and survival. For example, it could be a trait that increases fertility at the expense of life expectancy, or it could be a trait that consists in investing into survival at the expense of fertility. We start by considering a species living in a homogeneous environment, and we will see afterward how this approach can be generalized to account for the case of a plastic species living in a variable environment.

We consider a population consisting of *n* different genotypes. At time *t* the genotype *i* is in density *N*_*i*_(*t*) with *i* ∈ [1*, n*]. Total population density at *t* is therefore 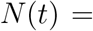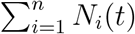. In what follows, for the sake of simplicity, we will sometimes drop the explicit dependency on time, but it always remains implicitly present. Each genotype is characterized by the value of a quantitative genetic trait *z*_*i*_ that potentially affects both mortality and fertility. In fact, the model is general in the sense that *z* could really represent *any* genetic trait.

Note that we deliberately choose not to consider an age-structured population. This will limit the scope of our results and we will point this out when this is the case. The advantage of such a simplification, however, is that it allows the reader to see the effect of extrinsic mortality in the most straightforward way possible. Related to this hypothesis, we also assume that the intensity of density-dependent competition is merely a function of total population density. In nature, different age classes or phenotypes may have different contributions to density-dependent competition –for example, adults versus immatures– but we ignore this source of complexity here.

1. We assume that an individual with a trait value *z* has a constant mortality rate over his lifetime, given by the function *d*(*z, N*) = *μ* + *m*(*z, N*) where *μ* is the extrinsic mortality, defined as the additive component of mortality that is independent of an individual’s age, condition or strategy (see Dańko et al. 2017), and *m*(*z, N*) represents the intrinsic mortality rate that depends both on the individual’s strategy, *z*, and also potentially on the intensity of competition in the population (hence on the total density *N*). Note that, in the sake of simplicity, we assume that density-dependent competition only affects the intrinsic mortality of individuals. Extrinsic mortality is supposed to be a fixed constant independent of demography.
2. We assume that an individual with trait *z* produces offspring at a constant fertility rate over his lifetime, given by the function *b*(*z, N*), which also potentially depends both on the individual’s strategy, and on the intensity of density-dependent competition. The form of the functions *b*(·) and *d*(·) need not be specified.

We consider a continuous-time model, as inspired by the epidemiological application of the Price equation developed by Day and Gandon (2005). The dynamics of the *n* genotypes in the population are given by the following system of differential equations

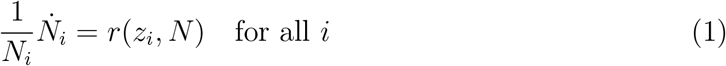

where *r*(*z, N*) = *b*(*z, N*) − *d*(*z, N*) is the net growth rate of each genotype. The frequency of each genotype being 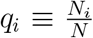, the dynamics of genotypic frequencies are given by the system

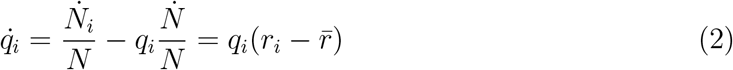

where the dot accents denote time derivatives (that is, for any variable *x*, 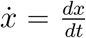), and 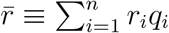 is the average *r*_*i*_ over the genotype distribution. The average value of *z* at time *t* is 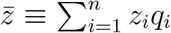, which therefore changes due to natural selection at a rate given by

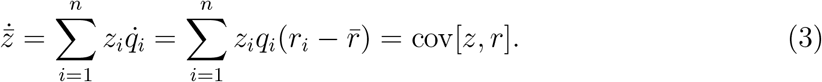

At each instant, the average value of *z* changes, under the effect of natural selection, at a rate given by the covariance between *z* and the instantaneous growth rate of genotypes, *r*. The trait *z* increases to the extent that is is associated with a larger growth rate, and vice versa. Equation 3 is a simple version of the so-called Price equation (Price, 1970).

To simplify the analysis, we now consider the case where the variability of *z* is small. In this case, we write the trait value of each allele as a deviation from the average trait value, i.e. 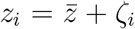. We then express the change in 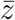 due to selection (Eq. 3), to the second order in *ζ*_*i*_. Assuming that all allelic effects scale as a common factor *ζ*, in the Appendix we show that, for small *ζ*, it simplifies into

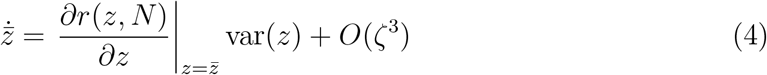

where 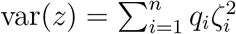 is the variance of *z* in the population, and *O*(*ζ*^3^) is at most a positive constant multiple of *ζ*^3^ as *ζ* → 0. In other words, to the second order, the change, due to selection, of the average value of *z* is simply proportional to the derivative of the growth rate *r* with respect to *z*. In what follows, we simply call this value the selection gradient on *z*, given by

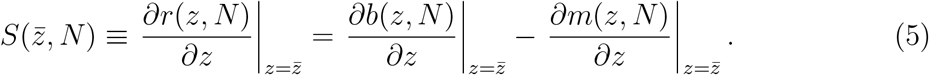

It is important to realize that this equation is not *dynamically sufficient*. It only shows the instantaneous effect of selection on *z* at any given time, provided one knows the mean trait value, 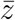, and the population density, *N*, at this time. This effect, however, does change over time, as 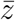 and *N* necessarily change, and equation 5 does not tell us how. As a result, it cannot be used to derive the dynamics of 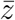 from an initial condition. It is nevertheless useful because it allows us to understand the effect of selection on a life history trait in a simple and general way. It does not imply any specific form of density-dependent competition –in fact it is valid even in the total absence of density-dependent competition– and it does not even assume that the population is at a demographic equilibrium. From this expression, our objective will be to understand how, and under what conditions, the selection gradient may depend on the extrinsic mortality of individuals.

### 3.2 Measuring the effect of selection in a variable environment

We can now clarify the extent to which the evolutionarily stable reaction norm in the case of a plastic species facing a variable environment can be inferred from the hard-wired evolutionarily stable strategies in different homogeneous environments, a question also discussed by Baldini (2015). To do so, we extend the above approach to consider the case where the environment is variable, either in space or time. For example, in some places or at certain times, extrinsic mortality is high, while it is low in others.

We assume that individuals are plastic in the expression of their life history trait *z*. In an environment of type *ε* they express a trait value *z*^*ε*^. For simplicity, we neglect (i) the cost of plasticity and (ii) the possible existence of a delay in the expression of plasticity. We simply assume that, in each type of environment, individuals are able to instantaneously express a life-history strategy specific to that environment, by modifying their allocation to survival and reproduction. Importantly, we also assume that the fecundity and mortality of individuals living in an environment of type *ε* only depend (i) on the properties of *ε*, and (ii) on the density *N*^*ε*^ of competitors in this very environment. That is, individuals are not affected by the environments in which they do not live (or only indirectly via *N*^*ε*^). Under these assumptions, our objective is to evaluate the direction of natural selection on each value of *z*^*ε*^.

An individual placed in an environment of type *ε* has fecundity *b*^*ε*^(*z*^*ε*^, *N*^*ε*^) and mortality *μ*^*ε*^ + *m*^*ε*^(*z*^*ε*^, *N*^*ε*^), where *N*^*ε*^ measures the intensity of density-dependent competition in the environment of type *ε*. As in the case of a homogeneous environment (Equation 5), the instantaneous direction of selection on *z*^*ε*^ is given by

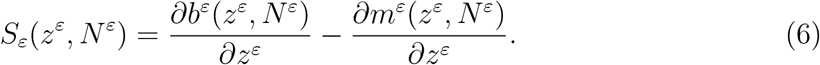

## 4 Results

### 4.1 Selection in a constant environment

The first thing we observe in equation 5 is that *μ* does not appear in the expression of *S*, hence that the selection gradient on the trait *z* cannot depend on extrinsic mortality (except through an effect on the other parameters in this equation; “indirect effect”, see below). In other words, the level of extrinsic mortality experienced by individuals, in itself, has no effect on selection for *any* trait, which is a standard result obtained by several classic papers (Abrams, 1993; Williams et al., 2006; Caswell, 2007) and see Moorad et al. (2019) for a review.

Here we find a discrepancy with Baldini (2015) who observes that, even in the absence of density-dependent regulation –thus without any effect of demography, what he calls “extrinsic mortality” does affect the evolutionarily stable life history, in a direction opposite to the standard view. This result, however, is actually a consequence of using a definition of extrinsic mortality that is inconsistent with previous works. Extrinsic mortality is standardly defined as the additive component of mortality that is independent of an individual’s age, condition or strategy, which amounts to a minimum mortality below which individuals cannot fall. By contrast, Baldini’s extrinsic mortality parameter is the individuals’ *maximum* mortality rate: the mortality rate that would be suffered by individuals who would not invest any resource at all in their survival. Thus defined, “extrinsic mortality” increases the marginal return on investment in survival, forcing individuals to invest more in survival and reproduce later, thus giving the impression of an effect opposite to the standard predictions. On the contrary, if defined properly, in the absence of density-dependent regulation extrinsic mortality merely has no effect on the evolution of life history traits.

However, the selection gradient on the trait *z* can depend *indirectly* on extrinsic mortality. Once again, it must not be forgotten that equation 5 is not dynamically sufficient, that is, it describes only the effects of extrinsic mortality over one time step on the selection gradient for given population density *N*, but not the effects that the expression of the trait *z* in previous time steps may have on *N*, and then on the selection gradient. The indirect effects are those not accounted by the dynamically insufficient result, in contrast to the direct effect of *z* for given *N*. All other things being equal, if extrinsic mortality increases then, *N*, is likely to decrease since individuals live shorter lives on average. In principle, it is therefore possible for extrinsic mortality to affect the evolution of life-history strategy indirectly, via its effect on *N* (since *N* appears in eq. 5). This effect could in principle be modelled under various scenarios, but we do not attempt to do so here. Our objective is simply to show that, if such an effect exists, then extrinsic mortality may have an effect on selection through this means.

#### 4.1.1 If pace of life and competition have independent effects, then extrinsic mortality plays no role in the evolution of pace of life

Let us first assume that the trait *z* and density-dependent competition affect fertility and mortality independently from each other, i.e., that they have additive effects. We write the fertility of a genotype *z* as *b*(*z, N*) = *b*_0_(*z*) − *αN*, and its mortality as *d*(*z, N*) = *μ* + *m*_0_(*z*) + *βN*, where *α* and *β* measure the intensity of the effect of competition, respectively, on fertility and survival.

In this case, the selection gradient on *z* is simply given by 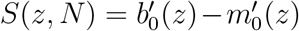 and is therefore independent of demography. This is logical since, in this situation, density-dependent competition affects all genotypes identically. As a result, extrinsic mortality cannot have any effect on the evolution of *z*. A particular case of this situation occurs in the total absence of density-dependent regulation (*α* = *β* = 0). In this case, extrinsic mortality also plays no role in the evolution of *z* (see also Moorad et al. 2019).

#### 4.1.2 If pace of life interacts with competition, then extrinsic mortality can affect its evolution

Let us now assume that the trait *z* and density-dependent competition interact with one another. We thus write the fertility of a genotype *z* as *b*(*z, N*) = *b*_0_(*z*) − *αN* − *γb*_0_(*z*)*N*, and its mortality as *d*(*z, N*) = *μ* + *m*_0_(*z*) + *βN* + *δm*_0_(*z*)*N*, where *γ* and *δ* measure the *interaction* effects between *z* and *N*, respectively on fertility and mortality. If *γ* > 0, then competition reduces more strongly the fertility of an individual who invests a lot in reproduction (if *γ* < 0, it is the opposite). If *δ* > 0, then competition increases more strongly the mortality of an individual who invests little in survival (if *δ* < 0, it is the opposite).

The direction of selection on *z* is given here by 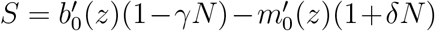. All other things being equal, selection is of course always positive on fertility and negative on mortality. What matters, however, is to measure the *relative* strength of selection on these two components. If selection is particularly strong on fertility, fast strategies are favoured. While, if selection is particularly strong on survival, slow strategies are favoured. Here the relative strength of selection on fertility and mortality is given by the ratio *F* = (1 − *γN*)(1 + *δN*)^−1^ which is therefore a measure of the strength of selection in favour of a fast pace of life. From this expression we find that, if it leads to a reduction of the intensity of density-dependent competition (lower *N*), extrinsic mortality can, in principle, affect the evolution of pace of life in two opposite directions.

1. Increased extrinsic mortality can favour a faster strategy with higher investment in fertility and lower investment in survival (higher *F*) in two cases:

- If competition reduces the benefit of investing in reproduction, by impacting especially strongly individuals with high fertility (*γ* > 0). This is typically the case if competition reduces the survival of immature offspring. Strictly speaking, our model does not capture this situation since we do not consider an age structure but, in an approximate way, an excess mortality of immatures results in a reduction of the number of offspring recruited in the next adult generation, per young produced. That is, it reduces the benefits of investing in reproduction.
- If competition increases especially strongly the mortality of individuals who invest little in their survival (*δ* > 0). This can happen if individuals with low somatic investment are particularly sensitive to competition, and have a mortality rate that increases sharply in the presence of competitors. This first situation seems reasonable and is likely to be frequent, for two reasons. First, the negative impact of competition on reproduction has good reasons to be stronger for individuals who invest a lot in reproduction (i.e. *γ* is likely to be positive). An individual who produces many offspring is likely to lose a large number of them to competition, in absolute terms, because she simply has more of them to lose. Second, the individuals who invest the most in reducing their mortality *in general* are likely to be well prepared to cope *in particular* with the effects of competition on mortality (i.e. *δ* is likely to be positive). For these two reasons, it is understandable that this effect –that a larger extrinsic mortality favours a faster strategy– is considered to be the most standard.
2. However, it is by no means the only possible effect. It is quite possible, under certain conditions, that the effect of an increased extrinsic mortality may go in the other direction, favouring slower strategies. This can occur in two cases:

- If competition has a particularly strong impact on the mortality of individuals who invest heavily in their survival (*δ* < 0).
- If competition increases the benefit of investing in reproduction by impacting less strongly individuals with high fertility (*γ* < 0). In practice, the first effect may occur, for example, if larger individuals, who survive better, are also more sensitive to resource depletion due to competition. The second effect, on the other hand, seems more of a theoretical possibility than a plausible situation in practice.

In summary, the key notion to remember is that extrinsic mortality can *only* affect the evolution of life history via its possible effect on competition and *by no other means*. To get an intuitive understanding of the effect of mortality implies to get an intuitive understanding of the effect of competition, nothing more. If a trait is an adaptation to intense competition that allows individuals to better thrive under this circumstance, for example by producing fewer but less fragile offspring, by investing in the ability to survive in a dense population, or by investing in the ability to compete for resources, then this trait is particularly favoured when extrinsic mortality is low. Conversely, if a trait is adapted to an environment with a low level of competition, but makes individuals especially sensitive to the presence of conspecifics, for example by producing numerous but fragile offspring, or by investing little in somatic features that allow to better compete, then this trait is favored when extrinsic mortality is high.

### 4.2 There is more to harshness than extrinsic mortality

We showed that extrinsic mortality can affect the pace of life only through its effect on density-dependent competition. But harsh environments can affect life-history evolution through other, more direct ways, as our model also shows.

To see this, let us consider the direction of selection on a trait *z* that reduces mortality at the expense of fecundity –i.e. a “slow” trait. Without loss of generality, suppose that *z* ∈ [0, 1] and that *z* has a linear effect on fecundity, with *b*(*z*, *N*) = *b*_0_ − *z* − *αN* + *γzN*, while mortality is given by *d*(*z*, *N*) = *μ* + *m*_0_(*z*) + *βN* + *δm*_0_(*z*)*N* where *m*_0_(*z*) is a decreasing function of *z*. The direction of selection on *z* is then given by 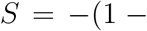 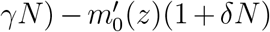. Beyond the effect of density-dependent competition (*γ* and *δ*), *S* also depends on 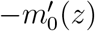, that is the marginal benefit from mortality reduction achieved by giving up one unit of fecundity. The higher this benefit (i.e. the more strongly negative is 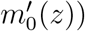 the more advantageous it is to invest in survival. Thus, even in the complete absence of density-dependent effects, it is quite possible that a feature of harsh environments related to the level and type of mortality risks may influence the life-history of individuals.

For example, one might speculate that harsh environments are not really characterized by a higher extrinsic mortality. Rather, harsh environments could be ones in which it is more costly to protect against mortality risks because they are more severe. In this case, 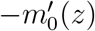 would be lower in harsh environments, making it less worthwhile to invest in survival, thereby favoring faster strategies, even in the absence of density-dependent effects.

One could speculate on the contrary that harsh environments are full of hazards but these are hazards against which it is fairly easy to protect oneself. In this case, the marginal benefit of investing in survival is greater and individuals may thus reduce their investment in fecundity. This is the effect captured in Baldini (2015), which actually considers maximum mortality despite calling it extrinsic mortality. On the other hand, it should be noted that, in this case, nothing can be said regarding the actual mortality of individuals at evolutionary stability without further specifying the shape of *m*_0_(*z*). Individuals will invest more in survival in harsh environments, but there is no certainty that they are able to fully compensate for the excess mortality.

More generally, this leads us to underline the fact that harsh and benign environments might differ not so much in terms of their extrinsic mortality (here *μ*) but in terms of the trade-off between mortality and fertility (here the function *m*_0_(.)). We will come back to this point in the Discussion.

### 4.3 Selection in a variable environment

In the case of a heterogeneous environment, from equation 6 we see that, in a local environment of any given type *ε*, extrinsic mortality does *not* directly affect the selection on *z*^*ε*^. But it affects it *indirectly* via its effect on the intensity of competition (measured by *N*^*ε*^). We can therefore qualitatively understand the effect of extrinsic mortality in a heterogeneous environment, as we did in a constant environment. If extrinsic mortality is particularly high in a given type of environment, this affects the direction of selection on the life history strategy expressed in this particular type of environment (i.e. it exerts a selective pressure on the reaction norm at this particular point), only to the extent that this higher mortality leads to a relaxation of competition in *this* type of environment. Conversely, if extrinsic mortality is particularly low in a given environment, it affects the direction of selection, only to the extent that this lower mortality leads to an enhanced competition in *this* type of environment.

We can therefore understand why the effect of selection in a variable environment may not be the same, quantitatively, as in a constant environment. When the environment is variable, the intensity of competition in a given type of environment does not depend solely on extrinsic mortality in *this* type of environment. If the migration rate from one environment to another is large, or if environment properties change rapidly through time, then population density in a given type of environment may also be affected by extrinsic mortality in other types. If extrinsic mortality is very high in a given environment, for instance, individuals living in this environment may still experience a relatively intense competition if they are surrounded –temporally or spatially– by other environments where mortality is low. As a consequence of this demographic coupling between environments, the difference between the life history strategies expressed by plastic individuals in two different environments is likely to be lower than the difference between the hard-wired strategies expressed by two populations evolving independently in different environments.

However, even though the magnitude of the effect of extrinsic mortality may not be the same in a variable than a constant environment, the *direction* of this effect is always the same, except under some unlikely scenarios. To see this, consider two environments that differ in terms of their level of extrinsic mortality. Say, extrinsic mortality is higher in environment of type *ε*_1_ than in environment of type *ε*_2_. We know from equation 6, that extrinsic mortality in environment *ε*_1_ may only affect the direction of selection in this environment via an effect on population density 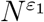. So, for the effect to reverse in the presence of environmental heterogeneity, population density would have to be *larger* in environment *ε*_2_ than in environment *ε*_1_, even though extrinsic mortality is higher. This seems almost impossible in principle since, if anything, extrinsic mortality has a negative effect on population density. However, it is conceivable that this could happen in specific scenarios where the migration rate of individuals is so high that the recruitment of dispersing individuals is higher than the recruitment of philopatric ones, or where the environment changes cyclically at a rate close to generation time since, in both cases, population density in one environment would depend more on the extrinsic mortality experienced by individuals in the *other* environment than in this one. But, apart from these specific scenarios, when extrinsic mortality decreases in a given environment, it may increase the intensity of competition less than in a fixed environment taken in isolation, but it does not reduce the intensity of competition in this environment; and conversely, if extrinsic mortality increases in a given environment, it does not increase the intensity of competition, even if it does not reduce it as much as in an environment taken in isolation. Therefore, even though the precise shape of the reaction norm cannot be predicted *quantitatively*, the direction of its slope can be reasonably well predicted *qualitatively* from the analysis of evolutionary stable strategies in constant environments.

Here as well, we observe a discrepancy with Baldini’s claims. But, as we pointed above, those claims are not based on the standard definition of extrinsic mortality. This matters here because the only quantitative result shown by Baldini in a heterogeneous environment (his Figure 1) concerns the case where density-dependence is absent, which is a situation where extrinsic mortality, if properly defined, has no effect on the evolution of life history anyway.

## 5 Discussion

In this article we have sought to clarify the effect of extrinsic mortality on the evolution of the ratio of investments in survival vs. reproduction, hereby referred to as the pace of life. Three intuitions on this point are common in the literature (see e.g. Ellis et al., 2009; Nettle, 2010; Belsky et al., 2010; Griskevicius et al., 2011; Frankenhuis et al., 2013; Mell et al., 2018): (I1) An environment with high extrinsic mortality always leads to the evolution of a fast pace of life (that is, a small investment in survival and a large investment in reproduction). (I2) This effect is due to the fact that, if mortality is high, individuals are likely to die before they have had time to reproduce. (I3) This effect is true both if the environment is constant and the pace of life a hard-wired strategy and if the environment is variable and the pace of life a plastic reaction norm.

In the present article, based on a simple model, we have sought to clarify the issue. We have also contrasted our results to those of Baldini (2015) and explained the discrepancies, but here we will emphasize the more positive messages from our model. We obtained the following results:

1. Strictly speaking, the three common intuitions above are false. In particular, intuition I2 is profoundly wrong. When extrinsic mortality does affect the evolution of pace of life, it is not for the intuitive reason that individuals have little time to reproduce, but for another reason related to the effect of extrinsic mortality on competition (see below). This point has already been known in the theoretical literature for a long time (Abrams, 1993; Williams et al., 2006; Caswell, 2007; and see Moorad et al., 2019 for a review).
2. Theoreticians are not hopeless, however, when it comes to predicting and explaining the effect of extrinsic mortality. With our simple model, we showed that it is possible to explain in a fairly intuitive and principled way the genuine effects of extrinsic mortality on pace of life. On this point our results are also in line with the theoretical literature (e.g. Abrams, 1993; Dańko et al., 2017), but the very simple nature of our model allows us show the effect of extrinsic mortality in the most straightforward way possible. Extrinsic mortality has no “direct” effect on the evolution of pace of life. However, it can affect it *indirectly* through its effect on the intensity of competition. Reducing extrinsic mortality always increases the intensity of competition because more individuals can be maintained in the environment. Hence, reducing extrinsic mortality favours slower strategies if these strategies allow to better thrive under intense competition and, on the contrary, faster strategies if these strategies do better under intense competition. Overall, all the traits that are adaptive under intense competition are favoured when extrinsic mortality is low and, conversely, all the traits that are adaptive when competition is relaxed are favoured when extrinsic mortality is high. This is undoubtedly different from intuition I2, but it can be explained to non-theoreticians in a simple manner. In particular, it must be stressed that whereas it is wrong to say that extrinsic mortality *always* favors a faster strategy, in many cases it actually does. This is notably the case in the very frequent situations where density-dependent regulation takes place via the fecundity of individuals or, with similar effects, through the mortality of juveniles. In this case, a reduction in extrinsic mortality leads to a reduction in the marginal return of investing in reproduction relative to survival, which does favour a slower strategy.
3. Intuition I3 is quantitatively wrong but still qualitatively true. Strictly speaking, models of homogeneous environments do not allow measurement of the effect of selection in variable environments. But the discrepancy between the two is only quantitative, not qualitative. Provided the cost of phenotypic plasticity is negligible, the *direction* of the effect of extrinsic mortality in a variable environment (but not the force of the effect) can be predicted from a homogeneous environment model. For example, if a particular model of homogeneous environment shows that the evolutionarily stable pace of life increases with extrinsic mortality, then we can deduce that the evolutionarily stable reaction norm in a variable envionment would also cause the pace of life to increase with extrinsic mortality.

One may now wonder about the consequences of this clarification. What can be said regarding the development of future research, both theoretical and empirical, on the application of life history theory to the understanding of human intraspecific variability? One first avenue of research would be to try and gain an empirical understanding of the nature of density-dependence in human populations, and more specifically in human populations whose ecology corresponds to the one in which we likely evolved. Another avenue of research, however, would be to shift away from an exclusive focus on extrinsic mortality, whose effects are limited and indirect, to adopt a broader perspective on what really constitutes harshness.

First, the key difference – or one of the key differences – between harsh and benign environments may well be a property related to mortality but not *extrinsic* mortality. Extrinsic mortality is very strictly defined as the additive component of mortality that is independent of individuals’ strategy. It does not represent the overall level of “danger” in the environment, as one might think, but the amount of danger against which the organism can do nothing, even if it were to invest an infinite amount of resources in protection. It can be argued that such an exogeneous mortality does not exist in real life, as one can always invest to guard against all sources of danger (Kaplan and Gangestad, 2015). A different view of mortality in harsh environments, probably a more accurate one, would therefore consist in saying that harsh environments are ones in which it costs more to reduce mortality because they involve more severe dangers. Investments in fecundity are thus comparatively more profitable there than investments in survival, since mortality cannot be significantly reduced at a reasonable cost anyway. In this case, unlike the case where it involves a higher *extrinsic* mortality, harshness does favour a faster strategy regardless of any frequency dependent effect (see section 4.2). The bottom line is that mortality is never really exogenous and is always the consequence of an interaction between environments and strategies. So, this interaction should actually be measured, and modelled, to understand the influence of environments on life histories.

We can then go even further and ask whether the relevant distinction between harsh and benign environments really has anything to do with mortality at all. Although it is reasonable to assume that mortality plays an important role when comparing regions of the world that differ with regard to, say, parasite pressure or predation rate, it is not necessarily the case when it comes to comparing the ecology of people with different socio-economic statuses within the same country. A large fraction of the higher mortality of people with low SES in developped countries is a *consequence* of a life-history strategy of investing less in survival and maintenance. This difference could thus perfectly well be caused by differences between these environments that have nothing to do with mortality in the first place.

In a recent unpublished work, Mell et al. (2019) obtained results along these lines in the case of the evolution of time horizon. Individuals exposed to deprivation have a propensity to discount future rewards more steeply than wealthier individuals. The standard explanation is that this is a consequence of higher extrinsic mortality. But Mell et al. (2019) suggest that this might actually rather be a consequence of wealth. Their idea is that rewards are not mere points to collect and put in a safe but assets that individuals can use to do useful things. Hence, delaying the collection of a reward creates an opportunity cost in the sense that during the waiting time, the benefits otherwise generated by the reward are lost. This cost is independent of mortality but it depends a lot on the individual’s current wealth. If someone can significantly improve their living condition –or prevent them from deteriorating– by using new resources wisely, they pay a high waiting cost. And vice versa, for someone whose condition is already plateauing anyway, waiting costs are low. It is therefore quite conceivable that an important property of the strategy of individuals, namely their temporal preferences, is unrelated to mortality. This could in fact be true more generally of other facets of people’s life history which could be determined by wealth, rather than by environmental features pertaining to mortality per se.

Lastly, life history is not just pace of life or time discounting. The life history strategy of an individual also includes their investment in various forms of biological capital, and it also includes the temporal variations of their investments over the lifetime, that is their “shape of life” and not only their pace of life (Baudisch, 2011; Jones et al., 2014). Understanding all these features requires still other models. Even more broadly, the concept of life history strategy is sometimes used to encompass also a variety of behaviours (e.g. religiosity, cooperation, etc.) that cannot be directly understood from models on pace of life or time discounting.

In conclusion, there are many difficulties and still some grey areas in the application of life history theory to human variability, but one must not throw out the baby with the bathwater by rejecting the entire theory on the grounds that it is not yet perfect. Understanding human variation based on resource allocation theory is a research program with an important goal and a solid rationale. We still need to better understand what harshness really means, better understand the nature of the adaptive responses it has selected for, and better prove empirically the very existence of phenotypic plastiticy, but these shortcomings should not lead us to reject, as a matter of principle, the approach as a whole.

## Acknowledgments

We thank Daniel Nettle, Willem Frankenhuis and three anonymous reviewers for their helpful suggestions and advice in writing this paper, and we thank Coralie Chevallier and Nicolas Baumard for the many discussions we had with them on this subject. xx was supported by ANR-17-EURE-0017 FrontCog and ANR-10-IDEX-0001-02 PSL.

## Appendix: Measuring the effect of selection to the second order

Here we derive the instantaneous rate of change in the mean value of the trait to the second order in trait variation. We write genotype *i*’s trait value as a deviation relative to the mean, i.e. 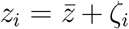, where 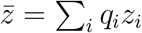 is the mean trait value, and therefore the mean deviation is zero by definition (Σ_*i*_ *q*_*i*_*ζ*_*i*_ = 0).

The instantaneous rate of change of the trait due to selection is

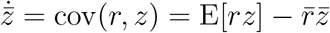

where E[*X*] stands for the expectation of *X* taken over the entire population.

We have

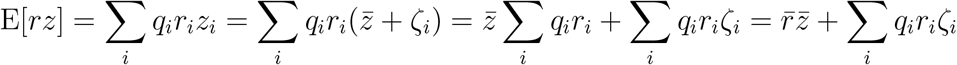

Hence this gives

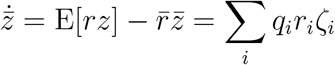

Quite generally for a function with a second derivative, for some 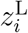 between 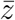 and *z*_*i*_, one can write 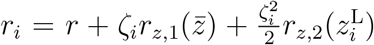, with *r*_*z,k*_(*x*) = *k*th derivative of *r* with respect to *z* evaluated for *z* = *x* (Taylor’s theorem with Lagrange form of the remainder). Then, using Σ_*i*_ *q*_*i*_*ζ*_*i*_ = 0,

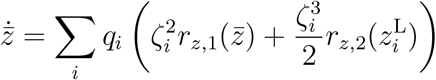

If the second derivative is bounded in absolute value over the range of all *z*_*i*_ values, then this reduces to

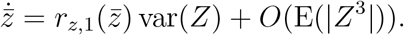

where we view each *ζ*_*i*_ as a realization of a random variable 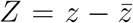. Assuming that all allelic effects scale as a common factor *ζ*, then var(*z*) = var(*Z*) = *O*(*ζ*^2^) dominates E(|*Z*^3^|) = *O*(*ζ*^3^) for small *ζ*, so that, to leading order in *ζ*, the instantaneous change in the mean trait value is given by the first derivative of *r* with respect to *z*(*r*_*z*,1_), multiplied by the variance of *z*:

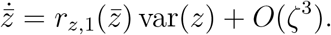

which is equation 4 of main text.

